# The inversion In(2L)t impacts complex, environmentally sensitive behaviors in *Drosophila melanogaster*

**DOI:** 10.1101/2025.09.17.676861

**Authors:** Benedict Adam Lenhart, Alan Olav Bergland

**Affiliations:** Department of Genome Science, University of Virginia, Charlottesville, VA 22904; Department of Biology, University of Virginia, Charlottesville, VA 22904

**Keywords:** Behavior, Startle Response, Foraging, Sleep

## Abstract

Genetic variation in behavioral traits allows organisms to respond and adapt to environmental challenges. Genetic variation in behavior is often affected by many genes and thus has a complex genetic basis. Inversions, the reorientation of genes along the chromosome, tightly link genetic variants together because they suppress recombination. Therefore, inversions are believed to have a major impact on phenotypic variation because they combine the effects of multiple genes, which can pleiotropically alter multiple aspects of behavior. This study investigates how the inversion In(2L)t, found in *Drosophila melanogaster* populations around the world, impacts different aspects of behavior in an environment-sensitive manner. We test the activity, foraging, and startle-induced behavior of flies with different In(2L)t genotypes across sex and temperatures. We observe that *Drosophila* homozygous for In(2L)t sleep less frequently, spend more time away from a food source, and have a longer duration of startle response. Additionally, the impacts of In(2L)t on aspects of behavior can be sex-specific and are largely consistent across temperatures. Taken together, our research demonstrates that inversions can regulate aspects of behavior, and suggests hypotheses explaining the distribution of In(2L)t across space and time.

## Introduction

Animals adjust their behavior in response to environmental change that occurs on multiple timescales. Long term changes like climate change influence habitat use and the timing of reproduction (Beever et al., 2017; Miller-Rushing et al., 2008), whereas short-term environmental changes like the day/night cycle affect foraging and activity levels (Ashby, 1972; Beauchamp, 2007). Some behavioral changes are cyclical, like the seasonal changes in migratory behavior (Chapman et al., 2015) or mating behavior (Riters & Stevenson, 2022). In general, behavioral variation is considered plastic in that a single individual modifies its behavior in response to environmental change (Snell-Rood, 2013). However, natural populations exhibit genetic variation in behavioral traits influencing both baseline behaviors (Fleury et al., 1995; Wong et al., 2019) and how behavior responds to environment factors (Flint, 2003; Niepoth & Bendesky, 2020).

Advancements in technology have improved our ability to quantify genetic and environmental effects on behavior (Pereira et al., 2020), and to do so at large scale (Schaefer & Claridge-Chang, 2012). For example, radio tags attached to mice track behavioral responses to variation in the nutritional environment and social cues (Peleh et al., 2019). Microscope recordings of *C. elegans* monitor responses to chemical and motor stimuli (Likitlersuang et al., 2012; Swierczek et al., 2011). In *Drosophila melanogaster*, automated methods have evolved from beam-break systems that inferred circadian activity and sleep (Chiu et al., 2010; Cichewicz & Hirsh, 2018; Pfeiffenberger et al., 2010) to video tracking software that quantifies speed (Faville et al., 2015) spatial positioning (Donelson et al., 2012), and responses to sensory stimuli (Werkhoven et al., 2019) often at scale (Elya et al., 2023; Werkhoven et al., 2019, 2021).

The ubiquity of genetic variation in phenotype (Mackay & Huang, 2018) and genotype (David & Capy, 1988), coupled with the ease of genetics and experimental study makes *D. melanogaster* a tractable model for studies into the natural genetic variation in behavior. One particularly interesting genetic variant is the large (10Mb) structural inversion, In(2L)t. This inversion is linked to elevated startle response duration (Mackay et al., 2012; Nunez et al., 2024) and foraging behavior (Lee et al., 2017). In(2L)t is found at intermediate frequencies in populations worldwide (Nunez et al., 2025; Stalker, 1980; van Delden & Kamping, 1989) and appears to mediate seasonal adaption in *D. melanogaster* as its frequency shifts seasonally (Nunez et al., 2024). Previous studies that report the behavioral impact of In(2L)t used the beam-break model or other methods with large-grain temporal sampling (Lee et al., 2017; Mackay et al., 2012; Nunez et al., 2024), providing the opportunity for better characterization of In(2L)t’s impact on behavior across a range of environments.

To explore how In(2L)t alters behavior in an environment-specific manner we employed automated behavioral tracking. We generated F1 offspring with different In(2L)t genotypes through controlled crosses and measured aspects of behavior of flies experiencing different temperatures. We found that In(2L)t is associated with increases in the duration of startle response, the time spent awake, and proximity to food. Furthermore, these inversion-driven differences often vary in intensity across sex, while being largely consistent across temperatures.

## Methods

### Fly stocks and husbandry

We measured behavior of F1s, generated through controlled crosses between inbred strains of the Drosophila Genetic Reference Panel (DGRP) (Mackay et al., 2012). By crossing five lines homozygous for In(2L)t (DGRP: 32,161, 348, 386, 837) and four lines homozygous for the “standard” gene arrangement (DGRP: 57, 189, 634, 853), we generated 8 unique F1 crosses homozygous for In(2L)t, 16 crosses heterozygous for In(2L)t, and 6 crosses homozygous for the standard genotype. Note, only a subset of possible crosses were used in this study. Flies were kept on standard media from Archon Scientific (Durham, NC; food is composed of 86% water, 0.574% agar, 6.30% cornmeal, 1.52% yeast, 4.65% molasses, 0.39% propionic acid, 0.15% methylparaben and 0.52% ethanol). Flies were reared from egg to adult in a 25°C incubator set to a 12H:12H light/dark schedule and 50% relative humidity. For each behavioral measurement, we used 2-5 day old non-virgin flies.

### Behavioral assays

We recorded the behavior of the F1 offspring using the *Drosophila* Arousal Threshold (DART) device (BFK Labs, Hertford, UK; Faville et al., 2015) set on top of an antivibration marble table. Flies were placed within plastic tubes with a 1.5% agar, 5% sucrose solution to prevent desiccation and starvation and a cotton plug to prevent asphyxiation. Before assays, we acclimated the flies in the DART overnight. The flies were kept in constant darkness, 50% relative humidity, and at three temperatures (20°, 25°, and 30°C). We recorded the basal activity, and induced activity following mechanical stimulus with a built-in motor. We performed this experiment in three replicate blocks, each time redoing the DGRP crosses and using the same behavioral quantification methods. The replicate experiments tested 414, 693, and 709 F1 individuals respectively. We measured male and female fly behavior at 25°C and then compared behavior of female flies at 20°C, 25°C and 30°C.

### Behavioral quantification

We used the DART MATLAB software (version 811f772; Faville et al., 2015) to track the motion of individual flies and quantify their behavior. We focused on three aspects of baseline behavior: duration of time spent sleeping (minutes per hour, m/h), speed (millimeters per second, mm/s), and the proportion of time a fly spends near the food. We assessed the sleep phenotype by identifying the sleep bouts defined as at least five minutes spent inactive for each fly, and then calculated the average minutes of time spent in sleep bout per hour. We assessed the baseline speed of the fly as the mean millimeters moved per minute in the hour prior to first stimulus.

We characterized fly startle response by measuring the duration of time with increased speed, and the magnitude of increased speed post stimulus. We first established individual fly baseline speed as the average speed in millimeters/second in the hour before the first stimulus, then estimated the duration of startle induced speed change as the time from the stimulus until the fly’s speed returned to its baseline speed. To estimate the magnitude of speed increase we calculated the difference in speed in the minute after the stimulus from the baseline speed.

To assess foraging behavior, we divided each tube into eight equal-length regions and calculated the proportion of time spent in the food-adjacent region compared to the other seven regions.

### Statistical analysis

We analyzed the impact of sex on behavioral traits using model comparison to identify the impact of inversion genotype, sex, as well as to test for interaction between inversion and sex. This analysis compared the observances of male and female flies exclusively at 25°C. We created the following models:

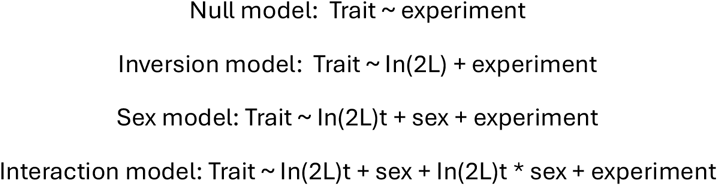

Where *ln(2L)t* is the fixed effect of inversion genotype, *sex* is the fixed effect of sex, and *experiment* is a random effect of the replicate experimental block. Mixed effect models were implemented in *lme4* (version 1.1-34, (Bates et al., 2015)). We compared the models using the *anova()* function from R version 4.3.1 to estimate the statistical significance of each of the fixed effects using likelihood ratio tests. To analyze the proportion of time flies spent near food, we first transformed proportions with an arcsin-square root transformation.

Next, we used model comparison to identify the impact of inversion genotype, temperature, as well as to test for gene-by-environment interaction between inversion and temperature. This analysis exclusively used data from female flies observed at 20°C, 25°C, and 30°C. We created the following models:

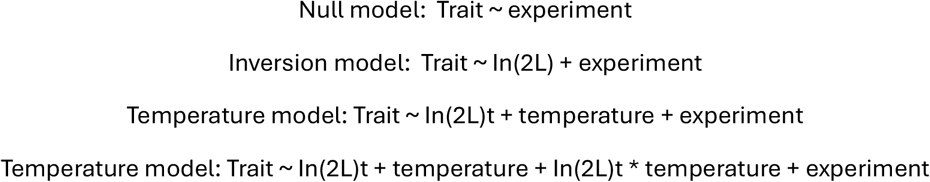

Where the *temperature* term is the fixed effect of temperature in Celsius of the environmental chamber, and the other terms are the same as above.

In both sets of models - either examining genotype and sex or genotype and temperature - if a significant effect was found using model comparison for the fixed effects of genotype or an interaction term including genotype, post-hoc pairwise Student’s t-tests were used to find differences between pairs of genotypes within sex or temperature.

## Results

*In(2L)t affects baseline levels of activity*. The amount of time flies spend sleeping is significantly affected by inversion genotype (Likelihood Ratio Test: *χ*^2^ = 51.29, df = 2, p = 7.3e-12; Table 1; **Fig. 1A**) and sex (*χ*^2^= 34.32, df = 1, p = 4.69e-9) when comparing the sexes at 25°C, with standard genotype flies and males sleeping longer on average. Female flies homozygous for the inversion sleep for less time than heterozygous flies (t-test: t = 3.93, df = 743 p = 9.28e-5), and heterozygote females sleep less than homozygous standard females (t = -4.65, df = 811, p = 3.85e-6; **Fig. 1A**). Male flies homozygous for the inversion also slept less than heterozygotes (t = -2.9, df = 106, p = 0.0045). Sex has a significant effect on time spent asleep (*χ*^2^ = 34.32, df = 1, p = 4.69e-09), but there is no interaction between sex and inversion genotype (*χ*^2^= 2.9, df = 2, p = 0.23).

**Figure 1:**
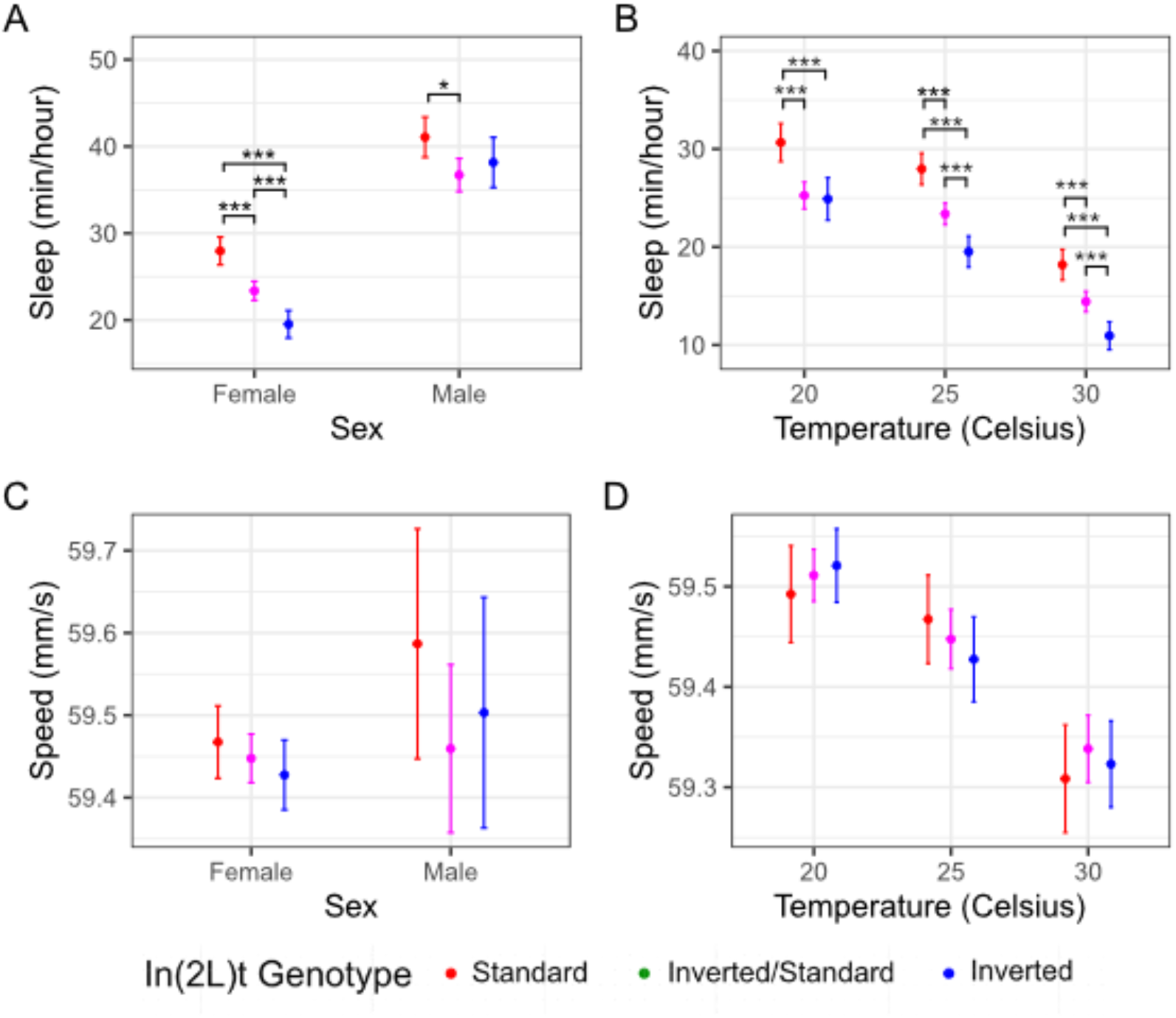
In(2L)t presence is associated with less time sleeping. A) Measurements of the minutes spent asleep per hour across sexes at 25C°, colored by the In(2L)t genotype. B) Comparison of time spent asleep over a range of temperatures for females. C) Baseline speed at 25°C measured in mm/s across sexes, colored by presence of In(2L)t. D) Same as in C, but comparing across temperatures. In all plots, the points represent the mean, and the error bars represent the 95% standard errors (* = p < 0.05, ** = p < 0.01, *** = p < 0.001, no * = p >= 0.05). Female measurements shown in panels A and B are redrawn in panels C and D, respectively, at 25°C.

**Table 1:** Summary statistics from model comparison of flies of both sexes. Comparison of Null, Inversion, Sex, and Interaction is given for each of the 6 traits reported on.

| Trait | Model | npar | AIC | logLik | Chisq | Df | Pr(>Chisq) |
| --- | --- | --- | --- | --- | --- | --- | --- |
| Sleep | Null Model | 3 | 15107 | -7550 |  |  |  |
| Sleep | Inversion Model | 5 | 15059 | -7525 | 51.29 | 2 | 7.30E-12 |
| Sleep | Sex Model | 6 | 15027 | -7507 | 34.32 | 1 | 4.69E-09 |
| Sleep | Interaction Model | 8 | 15028 | -7506 | 2.94 | 2 | 2.30E-01 |
| Base Speed | Null Model | 3 | 1851 | -922 |  |  |  |
| Base Speed | Inversion Model | 5 | 1850 | -920 | 4.71 | 2 | 9.51E-02 |
| Base Speed | Sex Model | 6 | 1851 | -920 | 1.02 | 1 | 3.13E-01 |
| Base Speed | Interaction Model | 8 | 1853 | -919 | 1.95 | 2 | 3.78E-01 |
| Startle Duration | Null Model | 3 | -2672 | 1339 |  |  |  |
| Startle Duration | Inversion Model | 5 | -2668 | 1339 | 0.41 | 2 | 8.15E-01 |
| Startle Duration | Sex Model | 6 | -2668 | 1340 | 2.05 | 1 | 1.52E-01 |
| Startle Duration | Interaction Model | 8 | -2665 | 1340 | 0.56 | 2 | 7.55E-01 |
| Startle Magnitude | Null Model | 3 | 3555 | -1775 |  |  |  |
| Startle Magnitude | Inversion Model | 5 | 3555 | -1772 | 4.73 | 2 | 9.40E-02 |
| Startle Magnitude | Sex Model | 6 | 3552 | -1770 | 5.20 | 1 | 2.26E-02 |
| Startle Magnitude | Interaction Model | 8 | 3555 | -1769 | 1.04 | 2 | 5.94E-01 |
| Prop_nearfood | Null Model | 3 | -516 | 261 |  |  |  |
| Prop_nearfood | Inversion Model | 5 | -517 | 263 | 4.83 | 2 | 8.95E-02 |
| Prop_nearfood | Sex Model | 6 | -515 | 263 | 0.17 | 1 | 6.77E-01 |
| Prop_nearfood | Interaction Model | 8 | -511 | 264 | 0.29 | 2 | 8.64E-01 |
| Prop_farfood | Null Model | 3 | -237 | 122 |  |  |  |
| Prop_farfood | Inversion Model | 5 | -269 | 140 | 35.89 | 2 | 1.61E-08 |
| Prop_farfood | Sex Model | 6 | -284 | 148 | 17.13 | 1 | 3.49E-05 |
| Prop_farfood | Interaction Model | 8 | -281 | 148 | 0.30 | 2 | 8.63E-01 |

Inversion genotype also significantly impacts sleep when considering females across temperatures (*χ*^2^= 84.19, df = 2, p < 2e-16; Table 2). At 20°C, heterozygous females slept as much as inverted homozygous females (t = -0.27, df = 606, p = 0.78: **Fig. 1B**), suggesting that the inverted allele is dominant, whereas at warmer temperatures the heterozygotes had an intermediate level of sleep, with standard flies sleeping longer than inverted flies (t = -6.74, df = 709, p = 3.35e-11). Temperature significantly impacts time spent asleep, with flies sleeping less at higher temperatures (*χ*^2^= 372.08, df = 1, p < 2e-16; Table 1). In general, differences in sleep behavior between genotypes were preserved across temperatures (**Fig. 1B**), and we did not observe gene-by-environment interaction (*χ*^2^= 4.43, df = 2, p = 0.11).

**Table 2:** Summary statistics from model comparison of female flies across temperatures. Comparison of Null, Inversion, Sex, and Interaction is given for each of the 6 traits reported on.

| Trait | Model | npar | AIC | logLik | Chisq | Df | Pr(>Chisq) |
| --- | --- | --- | --- | --- | --- | --- | --- |
| Sleep | Null Model | 3 | 37589.37 | -18791.68 |  |  |  |
| Sleep | Inversion Model | 5 | 37509.18 | -18749.59 | 84.19 | 2 | 5.24E-19 |
| Sleep | Temp Model | 6 | 37139.10 | -18563.55 | 372.08 | 1 | 6.60E-83 |
| Sleep | Interaction Model | 8 | 37138.67 | -18561.33 | 4.43 | 2 | 1.09E-01 |
| Base Speed | Null Model | 3 | 4173.44 | -2083.72 |  |  |  |
| Base Speed | Inversion Model | 5 | 4176.83 | -2083.42 | 0.61 | 2 | 7.36E-01 |
| Base Speed | Temp Model | 6 | 4058.91 | -2023.46 | 119.92 | 1 | 6.58E-28 |
| Base Speed | Interaction Model | 8 | 4062.51 | -2023.26 | 0.40 | 2 | 8.19E-01 |
| Startle Duration | Null Model | 3 | -6029.05 | 3017.52 |  |  |  |
| Startle Duration | Inversion Model | 5 | -6025.20 | 3017.60 | 0.15 | 2 | 9.28E-01 |
| Startle Duration | Temp Model | 6 | -6111.18 | 3061.59 | 87.98 | 1 | 6.62E-21 |
| Startle Duration | Interaction Model | 8 | -6108.56 | 3062.28 | 1.39 | 2 | 5.00E-01 |
| Startle Magnitude | Null Model | 3 | 8356.01 | -4175.01 |  |  |  |
| Startle Magnitude | Inversion Model | 5 | 8347.47 | -4168.74 | 12.54 | 2 | 1.89E-03 |
| Startle Magnitude | Temp Model | 6 | 8244.84 | -4116.42 | 104.63 | 1 | 1.47E-24 |
| Startle Magnitude | Interaction Model | 8 | 8240.66 | -4112.33 | 8.17 | 2 | 1.68E-02 |
| Prop_nearfood | Null Model | 3 | -1046.13 | 526.07 |  |  |  |
| Prop_nearfood | Inversion Model | 5 | -1060.68 | 535.34 | 18.55 | 2 | 9.39E-05 |
| Prop_nearfood | Temp Model | 6 | -1232.51 | 622.26 | 173.84 | 1 | 1.07E-39 |
| Prop_nearfood | Interaction Model | 8 | -1228.90 | 622.45 | 0.38 | 2 | 8.27E-01 |
| Prop_farfood | Null Model | 3 | -569.77 | 287.89 |  |  |  |
| Prop_farfood | Inversion Model | 5 | -639.37 | 324.68 | 73.59 | 2 | 1.05E-16 |
| Prop_farfood | Temp Model | 6 | -639.41 | 325.70 | 2.04 | 1 | 1.53E-01 |
| Prop_farfood | Interaction Model | 8 | -637.69 | 326.84 | 2.28 | 2 | 3.20E-01 |

We did not observe a significant difference in base speed between genotypes (*χ*^2^= 4.71, df = 2, p = 0.1; Table 1; **Fig. 1C**), sexes (*χ*^2^= 1.02, df = 1, p = 0.31), or any interaction between genotype and sex (*χ*^2^= 1.95, df = 2, p = 0.38) at 25°C. The inversion does not affect base speed in females across temperatures (*χ*^2^= 0.61, df = 2, p = 0.7365). However, there was a significant effect of temperature on speed with flies moving faster in warmer environments (*χ*^2^= 119.92, df = 2, p = <2e-16; Table2; **Fig. 1D**). There is no gene by environment interaction for speed (*χ*^2^= 0.40, df = 2, p = 0.82).

### lnversion genotype affects startle response duration and intensity

There is no effect of inversion genotype (*χ*^2^ = 0.41, df = 2, p = 0.82), sex (*χ*^2^ = 2.05, df = 1, p = 0.152; Table 1, **Fig. 2A**), nor inversion-sex interaction (*χ*^2^ = 0.56, df = 2, p = 0.76) on startle-induced changes in activity duration. Despite the lack of an overall effect, we identify that inverted female flies are startled for longer than standard females at 25°C (t = -2.23, df = 662, p = 0.0258). The effect of inversion genotype at 25°C is subtle, but consistent with previously published work. For instance, we reanalyzed existing datasets in Mackay et al., 2012 and found a significant increase in startle response duration for inverted females (Mackay et al., 2012), t = -3.38, df = 10, p = 0.007) and for inverted males ((Mackay et al., 2012), t = -2.86, df = 9, p = 0.02). When considering females across temperatures, there is no effect of genotype on the duration of startle-induced increase of speed (*χ*^2^ = 0.15, df = 2, p = 0.93; Table 2; **Fig. 2B**). Temperature does impact the startle response duration (*χ*^2^ = 88, df = 1, p < 2e-16), but there is no observed gene-by-environment interaction (*χ*^2^ = 1.39, df = 2, p = 0.50).

**Figure 2:**
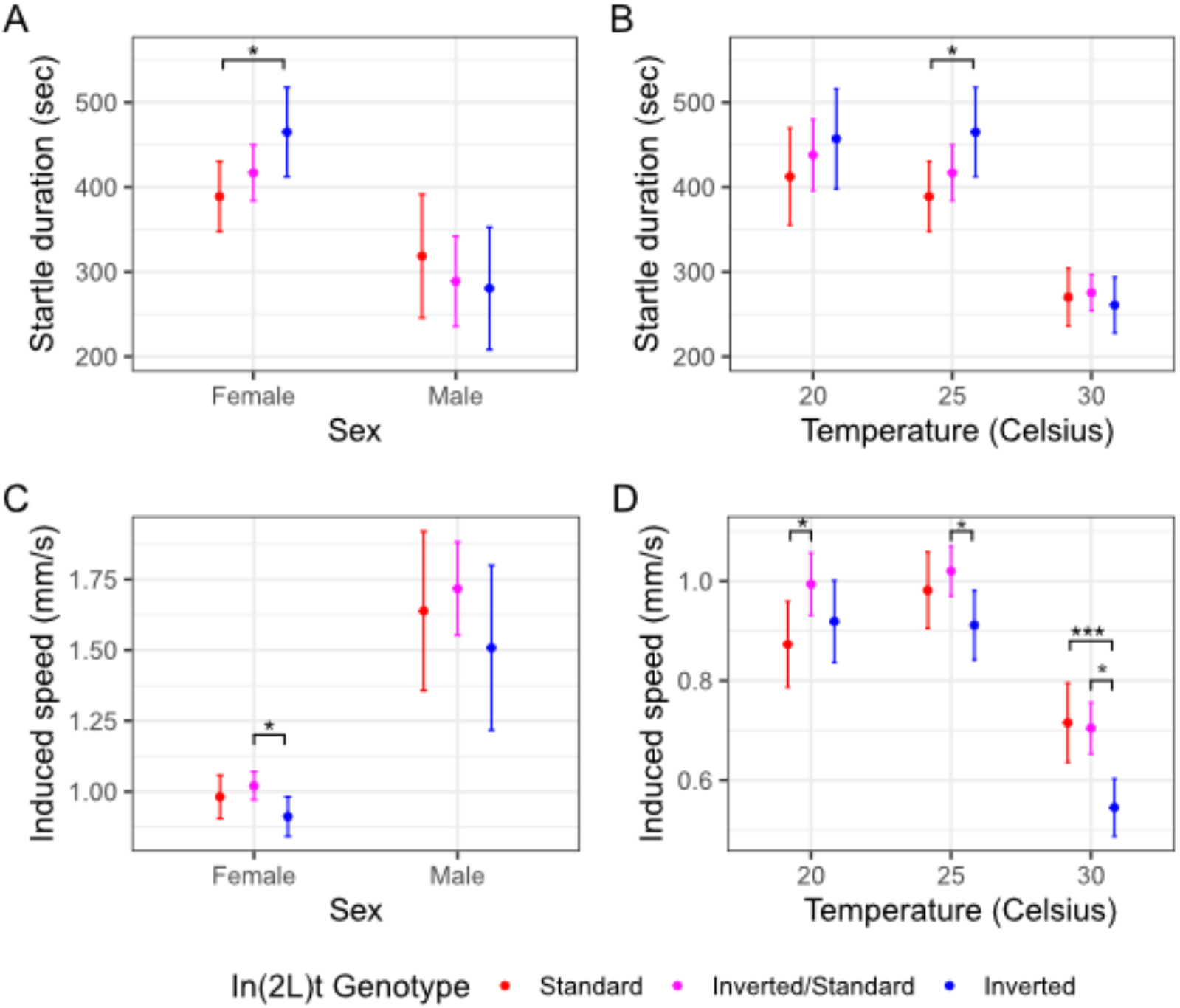
Inverted flies exhibit differences in startle induced speed. A) Startle response at 25°C measured as the duration of elevated response relative to baseline, colored by In(2L)t presence. B) Same as in A, but across temperatures. C) Induced activity at 25°C, defined as the difference in speed between startled state and baseline, again colored by In(2L)t genotype. D) Same as in C, but across temperatures. In all plots, the points represent the mean, and the error bars represent the 95% standard errors (* = p < 0.05, ** = p < 0.01, *** = p < 0.001, no * = p >= 0.05). Female measurements shown in panels A and B are redrawn in panels C and D, respectively, at 25°C.

Inversion genotype of the flies does not impact changes in speed following a startle (*χ*^2^ = 4.73, df = 2, p = 0.094; Table 1; **Fig. 2C**). The sex of the flies significantly impacts startle-induced speed (*χ*^2^ = 5.20, df = 1, p = 0.023), with male flies showing a larger response, and there is no observed genotype-sex interaction (X2 = 1.04, df = 2, p = 0.594). However, when considering female flies across temperatures, In(2L)t significantly impacts changes in speed following startle (*χ*^2^ = 12.54, df = 2, p = 0.0019; Table 2; **Fig. 2D**). Heterozygote females have higher induced speed than inverted homozygotes at 25°C (t = 2.5, df = 700, p = 0.013), while at 20°C heterozygote females have higher induced speed than standard homozygotes (t = -2.23, df = 662, p = 0.026; Fig. 2D). Inverted homozygotes have higher induced speed at 30°C than heterozygotes (t = -4.04, df = 790, p = 5.91e-5) or standard flies (t = 3.40, df = 612, p = 7.01e-4; **Fig. 2D**). Not only does temperature impact the fly speed following startle (*χ*^2^ = 104.63, df = 1, p = 0.0019), there is a gene-by-environment interaction observed in that the standard genotype decreases in its response compared to the other genotypes as temperature increases (*χ*^2^ = 81.7, df = 2, p = 0.017).

*In(2L)t is associated with location in relation to food*. When comparing the sexes at 25°C there is no effect of inversion genotype on proportion of time near food (*χ*^2^ = 4.83, df = 2, p = 0.09; Table 1; **Fig. 3A**), nor an effect of sex on proportion of time near food (*χ*^2^ = 0.07, df = 1, p = 0.68), nor a sex-genotype interaction (*χ*^2^ = 0.29, df = 2, p = 0.86). The amount of time spent near the food was strongly impacted by presence of In(2L)t while comparing females across temperatures (*χ*^2^ = 18.55, df = 2, p = 9.39e-05; Table 2; **Fig. 3B**). Female inverted flies spent less time near food than standard flies at 20°C (t = 1.98, df = 726.71, p = 0.048) and at 30°C (t = 3.09, df = 764.87, p = 0.0023). Temperature had a significant impact on the proportion of time flies spent near food (*χ*^2^ = 173.84, df = 1, p < 1.97e-16), though we did not observe any gene-by-environment interaction (*χ*^2^ = 0.38, df = 2, p = 0.83).

**Figure 3:**
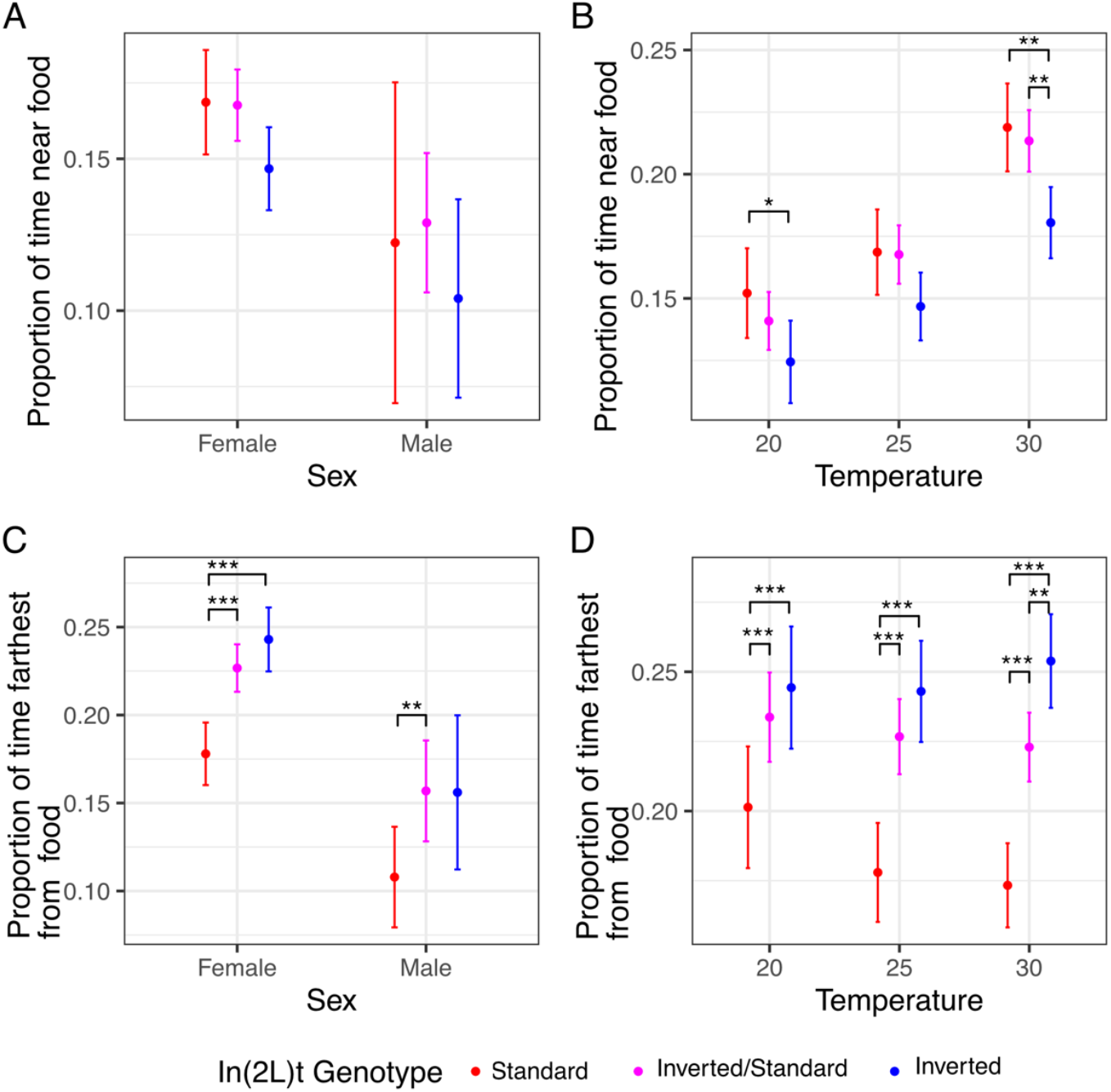
Inverted flies spend more time away from food source. A) Proportion of time spent in the region nearest food at 25°C within the DART experiment, colored by presence of In(2L)t. B) Same as in A, but across temperature. C) Proportion of time spent in the region farthest from food at 25°C within the DART experiment, colored by presence of In(2L)t. D) Same as in C, but across temperature. In all plots, the points represent the mean, and the error bars represent the 95% standard errors (* = p < 0.05, ** = p < 0.01, *** = p < 0.001, no * = p >= 0.05). Female measurements shown in panels A and C are redrawn in panel B and D, respectively.

We also examined the proportion of time flies spent on the end of their enclosure away from food. When observing flies at 25°C, we observe that inverted flies are away from food significantly more than standard flies (*χ*^2^ = 35.89, df = 2, p = 1.63e-08; Table 1; **Fig. 3C**). Female flies prefer the non-food end of the enclosure more than males (X2 = 17.13, df = 1, p = 3.59e-05), though there is no sex-inversion interaction (*χ*^2^ = 0.3, df = 2, p = 0.86). The preference for inverted female flies for the far end of the enclosure is observed across temperatures (*χ*^2^ = 73.59, df = 2, p = 1.05e-16; Table 2; **Fig. 3D**). Homozygous inverted flies spend more time away from food then standard at 20°C (t = -3.54, df = 641.73, p = 4.21e-04; **Fig. 3D**), 25°C (t = -6.09, df = 730.72, p = 1.88-09), and 30°C (t = -7.00, df = 646.21, p = 8.41e-12). There is no observed effect of temperature on time spent away from food (*χ*^2^ = 2.04, df = 1, p = 0.15), nor a temperature-inversion interaction (*χ*^2^ = 2.28, df = 2, p = 0.32).

Interestingly, Lee et al., 2017 also observed that In(2L)t is associated with variance in foraging within the DGRP (Lee et al., 2017). We reanalyzed their data and found that flies homozygous for In(2L)t were significantly worse at finding the food source, and subsequently starved sooner, than standard genotyped flies (t = 3.14, df =19, p = 0.005).

## Discussion

In this study, we use automated behavioral observation to characterize how the inversion In(2L)t impacts multiple aspects of behavior in an environment and sex specific manner. By quantifying motion and activity of *D. melanogaster* from a set of genetically diverse crosses, we show that presence of In(2L)t has an impact on time spent sleeping (**Fig. 1A, Fig.1B**), the duration and intensity of startle response (**Fig. 2**), and the amount of time spent near and away from food (**Fig. 3**). In(2L)t’s impact on behavior is mostly consistent across temperatures, though we do note evidence of gene by environment interaction between temperatures for startle response intensity (**Fig. 2D)**. With these findings, we emphasize the importance of inversions as a model for investigating the genetic variation in complex traits such as behavior. In(2L)t has a complex role in on behavior with impacts on baseline and induced activity, and contains genetic architecture thought to regulate aspects of behavior. ln general, homozygous In(2L)t flies spend less time sleeping than homozygous standard flies (**Fig. 1A**), and more often venturing out from the food source (**Fig. 3D**). Additionally, while In(2L)t flies may exhibit a reduced intensity of response to mechanical stimulus (**Fig. 2D**), they respond with elevated speed for longer than the standard genotype, at least at 25°C (**Fig. 2C**). The impact of In(2L)t on response to mechanical stimulus resembles In(2L)t’s impact on startle response observed in Mackay et al., 2012 (Mackay et al., 2012). There are several genes within In(2L)t that could explain the behavioral effects that we observe. The *foraging (for)* gene has a large impact on the range of travel while foraging (Nagle & Bell, 1987; Pereira & Sokolowski, 1993), and resides within In(2L)t. Indeed, previous work has identified that In(2L)t can explain part of the variance in foraging behavior (Lee et al., 2017), although that work link variation at *for* allele to foraging behavior. *Vglut* also resides within ln(2L)t. Found near the 5’ In(2L)t breakpoint, this gene regulates transport of glutamate, and is associated with sleep and startle response (Birgner et al., 2010; Hamasaka et al., 2007). Alleles for these genes and others within the inversion may explain the impact of In(2L)t genotype on different aspects of behavior, and may themselves be linked due to suppressed recombination (Dobzhansky & Epling, 1948; Kirkpatrick & Barton, 2006).

Season-specific regulation of behavior is essential for successful adaptation to new environments (Chapman et al., 2015; Riters & Stevenson, 2022). Strategies of inactivity versus activity are dependent upon the nutritional abundance (Wang et al., 2006), access to light (Welsh, 1938), and temperature stressors (Buchholz et al., 2019) that come with changing seasons. For example, circadian control of activity and sleep is closely regulated by light and temperature, two elements of the environment that change from summer to winter (Boothroyd et al., 2007). There is strong evidence that natural variation in sleep exists across different environments, with nighttime duration of inactivity increasing at lower latitudes (Svetec et al., 2015). Given that the length of sleep can be selected for (Harbison et al., 2017), this indicates selection could be acting on genetic variation regulating periods of inactivity in different latitudinal environments. Startle response behavior also shows evidence of adaptation to selective pressures, such as in the case of stickleback fish altering their startle response intensity depending on the temperature of their local environment (Guderley et al., 2001). Selection acts on many aspects of behavior in different directions across different seasons, but genetic variation in these traits must be preserved for this yearly adaptation to progress.

The effect of In(2L)t on behavior is largely consistent across temperatures, helping to define In(2L)t’s role in seasonal modification of traits. Inverted flies move more frequently than standard across temperatures (**Fig. 2B**), as well as more likely to be away from food (**Fig. 3D**). This consistent impact on behavior is relevant considering there is mounting evidence that In(2L)t is involved in mediation of seasonal adaptation. The frequency of In(2L)t changes cyclically across the year, becoming highest in the winter months (Machado et al., 2021; Nunez et al., 2024). This implies that the phenotypic variance explained by In(2L)t could be under directional selection within a given season. This argument is supported by the identification signals of selection for loci within the inversion (Nunez et al., 2024). A limited effect of the environment on genotypic effects on phenotype (lack of GxE), can lead to genotype-environment effects on fitness if different behavioral strategies are beneficial at different times of year, and can potentially drive the yearly fluctuation in In(2L)t frequency. flf higher activity is adaptive within colder months, then this could drive In(2L)t frequency higher, while high activity becoming maladaptive in warmer months could drive the In(2L)t frequency lower. The potential complex and shifting selection on the inversion can help explain both why In(2L)t frequency fluctuates yearly (Machado et al., 2021), and how this inversion continues to be identified at intermediate frequency within populations (Nunez et al., 2024; Stalker, 1980; van Delden & Kamping, 1989). Future study could build of the findings reported here by further identifying the genetic regions within In(2L)t that are impacting different aspects of behavior, and characterizing the fitness effects of high and low activity levels within seasonal environments.

## Data Availability

The DGRP lines used here are available from the Bloomington stock center in Indiana (https://bdsc.indiana.edu/). Inversion genotype tables for the lineages are all available from the DGRP website (http://dgrp2.gnets.ncsu.edu/data.html). The reanalyzed phenotype data from previous studies can be found from their original publications. All other data is available for download at (https://github.com/benedictlenhart/In-2l-t_Behavior)

## Acknowledgements

We gratefully acknowledge Richard Faville and the rest of the BFKlabs team in setting up and using the DART technology.

## References

Ashby, K. R. (1972). Patterns of daily activity in mammals. Mammal Review, 1(7-8), 171–185. 10.1111/j.1365-2907.1972.tb00088.x

Bates, D., Mächler, M., Bolker, B., & Walker, S. (2015). Fitting Linear Mixed-Effects Models Using lme4. Journal of Statistical Software, 67, 1–48. 10.18637/jss.v067.i01

Beauchamp, G. (2007). Exploring the role of vision in social foraging: What happens to group size, vigilance, spacing, aggression and habitat use in birds and mammals that forage at night? Biological Reviews, 82(3), 511–525. 10.1111/j.1469-185X.2007.00021.x

Beever, E. A., Hall, L. E., Varner, J., Loosen, A. E., Dunham, J. B., Gahl, M. K., Smith, F. A., & Lawler, J. J. (2017). Behavioral flexibility as a mechanism for coping with climate change. Frontiers in Ecology and the Environment, 15(6), 299–308. 10.1002/fee.1502

Birgner, C., Nordenankar, K., Lundblad, M., Mendez, J. A., Smith, C., le Grevès, M., Galter, D., Olson, L., Fredriksson, A., Trudeau, L.-E., Kullander, K., & Wallén-Mackenzie, A. (2010). VGLUT2 in dopamine neurons is required for psychostimulant-induced behavioral activation. Proceedings of the National Academy of Sciences, 107(1), 389–394. 10.1073/pnas.0910986107

Boothroyd, C. E., Wijnen, H., Naef, F., Saez, L., & Young, M. W. (2007). Integration of Light and Temperature in the Regulation of Circadian Gene Expression in Drosophila. PLOS Genetics, 3(4), e54. 10.1371/journal.pgen.0030054

Buchholz, R., Banusiewicz, J. D., Burgess, S., Crocker-Buta, S., Eveland, L., & Fuller, L. (2019). Behavioural research priorities for the study of animal response to climate change. Animal Behaviour, 150, 127–137. 10.1016/j.anbehav.2019.02.005

Chapman, J. W., Reynolds, D. R., & Wilson, K. (2015). Long-range seasonal migration in insects: Mechanisms, evolutionary drivers and ecological consequences. Ecology Letters, 18(3), 287–302. 10.1111/ele.12407

Chiu, J. C., Low, K. H., Pike, D. H., Yildirim, E., & Edery, I. (2010). Assaying Locomotor Activity to Study Circadian Rhythms and Sleep Parameters in Drosophila. Journal of Visualized Experiments : JoVE, 43, 2157. 10.3791/2157

Cichewicz, K., & Hirsh, J. (2018). ShinyR-DAM: A program analyzing Drosophila activity, sleep and circadian rhythms. Communications Biology, 1(1), 1–5. 10.1038/s42003-018-0031-9

David, J. R., & Capy, P. (1988). Genetic variation of Drosophila melanogaster natural populations. Trends in Genetics, 4(4), 106–111. 10.1016/0168-9525(88)90098-4

Dobzhansky, T., & Epling, C. (1948). The Suppression of Crossing Over in Inversion Heterozygotes of Drosophila Pseudoobscura. Proceedings of the National Academy of Sciences of the United States of America, 34(4), 137–141. 10.1073/pnas.34.4.137

Donelson, N., Kim, E. Z., Slawson, J. B., Vecsey, C. G., Huber, R., & Griffith, L. C. (2012). High-Resolution Positional Tracking for Long-Term Analysis of Drosophila Sleep and Locomotion Using the “Tracker” Program. PLOS ONE, 7(5), e37250. 10.1371/journal.pone.0037250

Elya, C., Lavrentovich, D., Lee, E., Pasadyn, C., Duval, J., Basak, M., Saykina, V., & de Bivort, B. (2023). Neural mechanisms of parasite-induced summiting behavior in ‘zombie’Drosophila. Elife, 12, e85410.

Faville, R., Kottler, B., Goodhill, G. J., Shaw, P. J., & van Swinderen, B. (2015). How deeply does your mutant sleep? Probing arousal to better understand sleep defects in Drosophila. Scientific Reports, 5(1), 8454. 10.1038/srep08454

Fleury, F., Allemand, R., Fouillet, P., & Boulétreau, M. (1995). Genetic variation in locomotor activity rhythm among populations ofLeptopilina heterotoma (Hymenoptera: Eucoilidae), a larval parasitoid ofDrosophila species. Behavior Genetics, 25(1), 81–89. 10.1007/BF02197245

Flint, J. (2003). Analysis of quantitative trait loci that influence animal behavior. Journal of Neurobiology, 54(1), 46–77. 10.1002/neu.10161

Guderley, H., Leroy, P. H., & Gagné, A. (2001). Thermal Acclimation, Growth, and Burst Swimming of Threespine Stickleback: Enzymatic Correlates and Influence of Photoperiod. Physiological and Biochemical Zoology, 74(1), 66–74. 10.1086/319313

Hamasaka, Y., Rieger, D., Parmentier, M.-L., Grau, Y., Helfrich-Förster, C., & Nässel, D. R. (2007). Glutamate and its metabotropic receptor in Drosophila clock neuron circuits. Journal of Comparative Neurology, 505(1), 32–45. 10.1002/cne.21471

Harbison, S. T., Negron, Y. L. S., Hansen, N. F., & Lobell, A. S. (2017). Selection for long and short sleep duration in Drosophila melanogaster reveals the complex genetic network underlying natural variation in sleep. PLOS Genetics, 13(12), e1007098. 10.1371/journal.pgen.1007098

Kirkpatrick, M., & Barton, N. (2006). Chromosome inversions, local adaptation and speciation. Genetics, 173(1), 419–434. 10.1534/genetics.105.047985

Lee, Y. C. G., Yang, Q., Chi, W., Turkson, S. A., Du, W. A., Kemkemer, C., Zeng, Z.-B., Long, M., & Zhuang, X. (2017). Genetic Architecture of Natural Variation Underlying Adult Foraging Behavior That Is Essential for Survival of Drosophila melanogaster. Genome Biology and Evolution, 9(5), 1357–1369. 10.1093/gbe/evx089

Likitlersuang, J., Stephens, G., Palanski, K., & Ryu, W. S. (2012). C. elegans Tracking and Behavioral Measurement. Journal of Visualized Experiments : JoVE, 69, 4094. 10.3791/4094

Machado, H. E., Bergland, A. O., Taylor, R., Tilk, S., Behrman, E., Dyer, K., Fabian, D. K., Flatt, T., Gonzalez, J., Karasov, T. L., Kim, B., Kozeretska, I., Lazzaro, B. P., Merritt, T. J., Pool, J. E., O’Brien, K., Rajpurohit, S., Roy, P. R., Schaeffer, S. W., . Petrov, D. A. (2021). Broad geographic sampling reveals the shared basis and environmental correlates of seasonal adaptation in Drosophila. eLife, 10, e67577. 10.7554/eLife.67577

Mackay, T. F. C., & Huang, W. (2018). Charting the genotype-phenotype map: Lessons from the Drosophila melanogaster Genetic Reference Panel. WIREs Developmental Biology, 7(1), e289. 10.1002/wdev.289

Mackay, T. F. C., Richards, S., Stone, E. A., Barbadilla, A., Ayroles, J. F., Zhu, D., Casillas, S., Han, Y., Magwire, M. M., Cridland, J. M., Richardson, M. F., Anholt, R. R. H., Barrón, M., Bess, C., Blankenburg, K. P., Carbone, M. A., Castellano, D., Chaboub, L., Duncan, L., . Gibbs, R. A. (2012). The Drosophila melanogaster Genetic Reference Panel. Nature, 482(7384), 173–178. 10.1038/nature10811

Miller-Rushing, A. J., Lloyd-Evans, T. L., Primack, R. B., & Satzinger, P. (2008). Bird migration times, climate change, and changing population sizes. Global Change Biology, 14(9), 1959–1972. 10.1111/j.1365-2486.2008.01619.x

Nagle, K. J., & Bell, W. J. (1987). Genetic control of the search tactic ofDrosophila melanogaster: An ethometric analysis ofrover/sitter traits in adult flies. Behavior Genetics, 17(4), 385–408. 10.1007/BF01068138

Niepoth, N., & Bendesky, A. (2020). How Natural Genetic Variation Shapes Behavior. Annual Review of Genomics and Human Genetics, 21(Volume 21, 2020), 437–463. 10.1146/annurev-genom-111219-080427

Nunez, J. C. B., Coronado-Zamora, M., Gautier, M., Kapun, M., Steindl, S., Ometto, L., Hoedjes, K. M., Beets, J., Wiberg, R. A. W., Mazzeo, G. R., Bass, D. J., Radionov, D., Kozeretska, I., Zinchenko, M., Protsenko, O., Serga, S., Amor-Jimenez, C., Casillas, S., Sanchez-Gracia, A., . Gonzalez, J. (2025). Footprints of worldwide adaptation in structured populations of D. melanogaster through the expanded DEST 2.0 genomic resource (p. 2024.11.10.622744). bioRxiv. 10.1101/2024.11.10.622744

Nunez, J. C. B., Lenhart, B. A., Bangerter, A., Murray, C. S., Mazzeo, G. R., Yu, Y., Nystrom, T. L., Tern, C., Erickson, P. A., & Bergland, A. O. (2024). A cosmopolitan inversion facilitates seasonal adaptation in overwintering Drosophila. Genetics, iyad207. 10.1093/genetics/iyad207

Peleh, T., Bai, X., Kas, M. J. H., & Hengerer, B. (2019). RFID-supported video tracking for automated analysis of social behaviour in groups of mice. Journal of Neuroscience Methods, 325, 108323. 10.1016/j.jneumeth.2019.108323

Pereira, H. S., & Sokolowski, M. B. (1993). Mutations in the larval foraging gene affect adult locomotory behavior after feeding in Drosophila melanogaster. Proceedings of the National Academy of Sciences, 90(11), 5044–5046. 10.1073/pnas.90.11.5044

Pereira Shaevitz, J.W., & Murthy, M. (2020). Quantifying behavior to understand the brain. Nature Neuroscience, 23(12), 1537–1549. 10.1038/s41593-020-00734-z

Pfeiffenberger, C., Lear, B. C., Keegan, K. P., & Allada, R. (2010). Locomotor Activity Level Monitoring Using the Drosophila Activity Monitoring (DAM) System. Cold Spring Harbor Protocols, 2010(11), pdb.prot5518. 10.1101/pdb.prot5518

Riters, L. V., & Stevenson, S. A. (2022). Using seasonality and birdsong to understand mechanisms underlying context-appropriate shifts in social motivation and reward. Hormones and Behavior, 142, 105156. 10.1016/j.yhbeh.2022.105156

Schaefer, A. T., & Claridge-Chang, A. (2012). The surveillance state of behavioral automation. Current Opinion in Neurobiology, 22(1), 170–176. 10.1016/j.conb.2011.11.004

Snell-Rood, E. C. (2013). An overview of the evolutionary causes and consequences of behavioural plasticity. Animal Behaviour, 85(5), 1004–1011. 10.1016/j.anbehav.2012.12.031

Stalker, H. D. (1980). CHROMOSOME STUDIES IN WILD POPULATIONS OF DROSOPHILA MELANOGASTER. II. RELATIONSHIP OF INVERSION FREQUENCIES TO LATITUDE, SEASON, WING-LOADING AND FLIGHT ACTIVITY. Genetics, 95(1), 211–223. 10.1093/genetics/95.1.211

Svetec, N., Zhao, L., Saelao, P., Chiu, J. C., & Begun, D. J. (2015). Evidence that natural selection maintains genetic variation for sleep in Drosophila melanogaster. BMC Evolutionary Biology, 15(1), 41. 10.1186/s12862-015-0316-2

Swierczek, N. A., Giles, A. C., Rankin, C. H., & Kerr, R. A. (2011). High-throughput behavioral analysis in C. elegans. Nature Methods, 8(7), 592–598. 10.1038/nmeth.1625

van Delden, W., & Kamping, A. (1989). THE ASSOCIATION BETWEEN THE POLYMORPHISMS AT THE Adh AND aGpdh LOCI AND THE In(2L)t INVERSION IN DROSOPHILA MELANOGASTER IN RELATION TO TEMPERATURE. Evolution, 43(4), 775–793. 10.1111/j.1558-5646.1989.tb05176.x

Wang, T., Hung, C. C. Y., & Randall, D. J. (2006). THE COMPARATIVE PHYSIOLOGY OF FOOD DEPRIVATION: From Feast to Famine. Annual Review of Physiology, 68(Volume 68, 2006), 223–251. 10.1146/annurev.physiol.68.040104.105739

Welsh, J. H. (1938). Diurnal Rhythms. The Quarterly Review of Biology, 13(2), 123–139. 10.1086/394554

Werkhoven, Z., Bravin, A., Skutt-Kakaria, K., Reimers, P., Pallares, L. F., Ayroles, J., & de Bivort, B. L. (2021). The structure of behavioral variation within a genotype. eLife, 10, e64988. 10.7554/eLife.64988

Werkhoven, Z., Rohrsen, C., Qin, C., Brembs, B., & Bivort, B. de. (2019). MARGO (Massively Automated Real-time GUI for Object-tracking), a platform for high-throughput ethology. PLOS ONE, 14(11), e0224243. 10.1371/journal.pone.0224243

Wong, W.-R., Brugman, K. I., Maher, S., Oh, J. Y., Howe, K., Kato, M., & Sternberg, P. W. (2019). Autism-associated missense genetic variants impact locomotion and neurodevelopment in Caenorhabditis elegans. Human Molecular Genetics, 28(13), 2271–2281. 10.1093/hmg/ddz051

